# Using EEG to Detect Lapses in Sustained Attention to Moving Stimuli

**DOI:** 10.1101/2025.07.10.663816

**Authors:** Benjamin G. Lowe, Alexandra Woolgar, Sophie Smit, Anina N. Rich

## Abstract

Sustaining attention is effortful but crucial for daily life. Despite this, attentional lapses are common and can have fatal consequences (e.g., when driving). The spontaneous nature of these lapses make studying their underlying phenomena elusive. As such, methods capable of determining when lapses have occurred may be fruitful research tools, with the potential to save lives if implemented within real world settings. Here, we capitalised on a recent hierarchical classification method, which uses multivariate decoding to index how well human observers sustain their attention within a dynamic visual environment. We asked whether this method could be used to anticipate behavioural errors based on neural activity measured with electroencephalography (EEG). We first decoded patterns of EEG activity that systematically correlated with critical aspects of a Multiple Object Monitoring (MOM) task. The extent to which we could decode this information depended on whether a stimulus was relevant for behaviour, which was lower before participants failed to detect (or ‘missed’) target stimuli, presumably due to attentional lapses. Here, we exploited this drop in neural decodability to predict whether errors were about to occur on each trial. The results form a foundation for sensitive and specific methods to objectively detect lapses in sustained attention based on patterns of brain activity.

## Introduction

Sustaining our attention on one task for a prolonged period is effortful. Because of this, momentary lapses in sustained attention are common and often disrupt goal-directed behaviour. While the consequences of these could be negligible (e.g., losing your place whilst reading a paper), there are many scenarios where lapses can be extremely costly or dangerous (e.g., when operating a vehicle). Thus, it is important to understand both the behavioural and neural underpinnings of attentional lapses.

By virtue of being spontaneous and internally driven, lapses in sustained attention are difficult to manipulate and study within experimental settings. Because of this, researchers often rely on participants self-reporting when lapses occur (Rasmussen et al., 2024; Smallwood et al., 2004; Unsworth & Robison, 2016), or indirectly infer this (post hoc) from poor task performance (deBettencourt et al., 2018; Karimi-Rouzbahani et al., 2021; Ladouce et al., 2024; O’Connell et al., 2009; Smallwood et al., 2004; Thomson, Smilek, et al., 2015). Ideally, we would have tools capable of objectively identifying when lapses occur in real time, allowing us to further study phenomena like lapse duration, termination, and recovery.

Using electroencephalography (EEG), prior studies have shown that trial-wise univariate measures of brain activity (e.g., posterior alpha and steady-state visually evoked potentials) can track when attentional lapses and/or fluctuations occur (e.g., Chinchani et al., 2022; Mazaheri et al., 2009; O’Connell et al., 2009; Renton et al., 2021). Because these measures are assumed to reflect the fidelity of stimulus encoding (Ergenoglu et al., 2004; Griffiths et al., 2019; Norcia et al., 2015), it is reasonable to assume that attentional lapses must attenuate the neural representation of *task-critical information*, which can be measured using multivariate decoding techniques (Grootswagers et al., 2017; Robinson et al., 2023). In other words, the trial-by-trial decodability of stimulus features should serve as a direct index of when participants are off-task or experiencing an attentional lapse.

Recent magnetoencephalography (MEG) work by Karimi-Rouzbahani et al. (2021) has given credence to this idea. Briefly, the authors trained linear classifiers to decode critical aspects of a sustained attention task that employed dynamic and overlapping stimuli, and then used the resulting decodability as the basis of a second-order classifier to predict which trials ended in behavioural errors. The amount of related information detectable within neural activity was correlated with both attention-related experimental manipulations (higher for task-relevant than task-irrelevant information) and whether participants were about to make errors (higher on ‘hits’ than ‘misses’). Crucially, these neural patterns were predictive of whether participants were about to miss targets with ∼65% accuracy for nearly an entire second before behavioural responses were required. Under the assumption that missed targets were predominantly caused by spontaneous lapses in sustained attention, this work suggests that multivariate representations of task-critical stimulus features may serve as a fruitful approach for tracking whether participants are on task.

This pre-registered study aimed to replicate and further extend Karimi-Rouzbahani et al.’s (2021) findings—this time using EEG instead of MEG, due to the former being significantly more affordable to setup and use. To preview our findings, we were able to demonstrate clear effects of sustained attention from task-critical patterns of neural activity whilst participants completed a dynamic Multiple Object Monitoring (MOM) task (Figure 1A). Crucially, we could statistically predict whether participants were about to miss targets, albeit with weaker classification accuracy than the initial MEG findings. This provides a proof-of-concept and foundation for future work developing sensitive and accessible methods for detecting attentional lapses in complex and dynamic environments from neural data.

**Figure 1.**
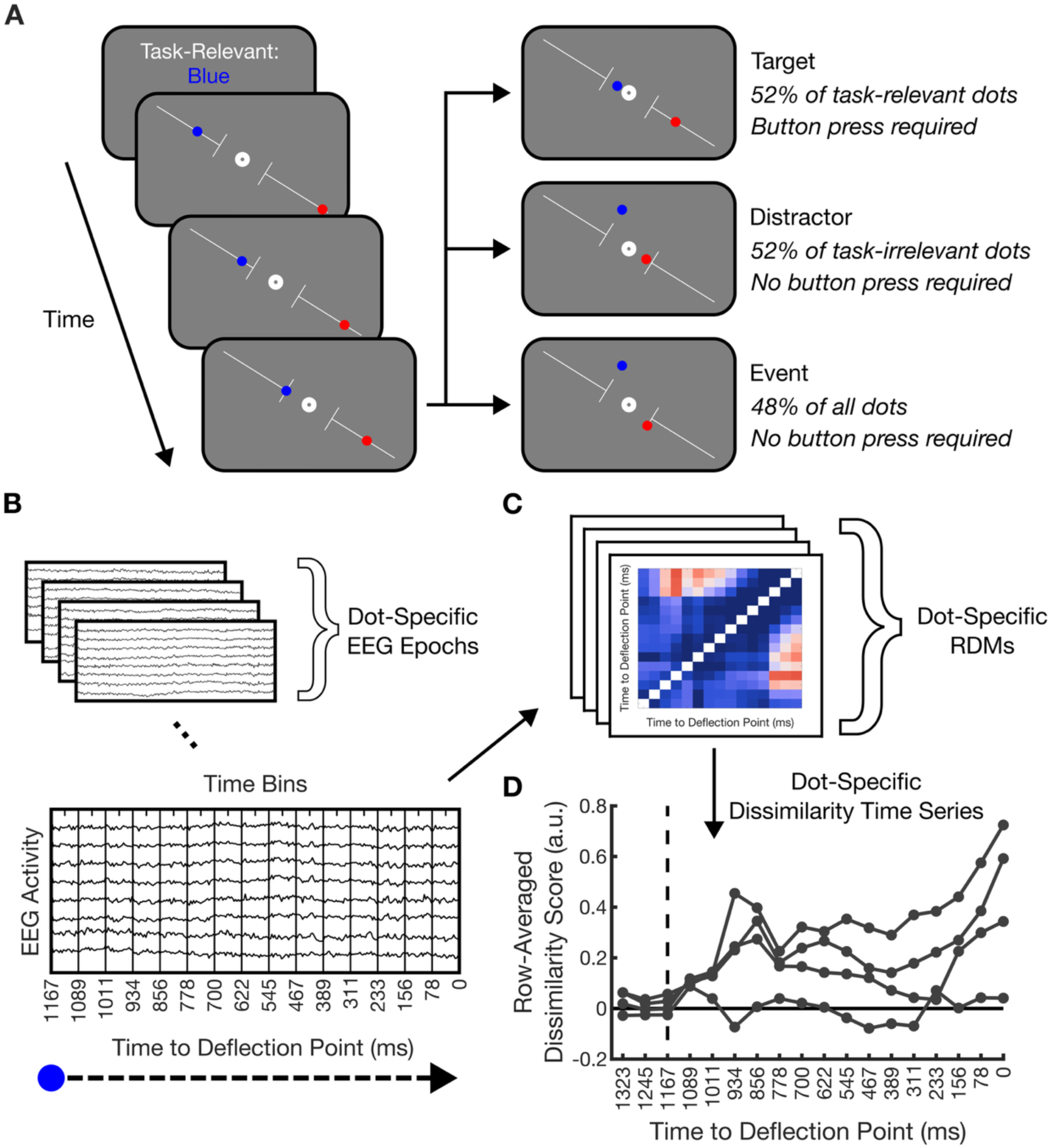
Multiple Object Monitoring (MOM) task and analysis pipeline. **A)** Schematic of the MOM task (video demonstration: https://osf.io/69ptm/). Each task block began with the participants being cued towards a task-relevant dot colour (here blue; task-irrelevant dot colour is red) followed by ∼2 min of a continuous task period. During the task periods, 100 dots onset (50 from the top-left and 50 from the bottom-right of the display, intermingled) and smoothly moved along visible trajectories towards the obstacle at fixation. Forty-eight percent of dots automatically deflected away from ‘colliding’ with the central obstacle once reaching the deflection point at the end of the trajectory (events). The remaining dots required manual deflection via a button press if they were the task-relevant colour (targets). Task-irrelevant dots that continued past the deflection points were to be ignored (distractors). **B)** The continuous EEG data were epoched relative to the onset of each dot. These epochs were further segmented into 77.8 ms time bins. **C)** Each dot-specific epoch was converted into a representational dissimilarity matrix (RDM) that characterised how dissimilar the multivariate patterns of EEG were between time bins. **D)** Dot-specific RDMs were converted into pseudo time-series data by averaging across dissimilarity scores from pairwise comparisons that shared a common time bin (i.e., averaging across the rows within each column). Steps **C)** and **D)** were also conducted on three pre-onset time bins to provide a benchmark of dissimilarity scores that could be achieved by chance.

## Methods

### Pre-registration Information and Data Availability

We pre-registered this study on the Open Science Framework before any data were analysed (https://osf.io/cu87z). Minor deviations from our pre-registration are detailed and justified within the Supplementary Material. All written code and analysed data will be made publicly available upon publication.

### Participants

Thirty healthy human participants were recruited through Macquarie University’s online participation pool (SONA) and the local community. Inclusion criteria required (corrected to) normal vision, normal hearing, fluency in speaking and reading English, and no significant history of neurological injury—all indicated via self-report. All participants gave informed written consent and were compensated with course credit or $40 (AUD). The study was approved by Macquarie University’s Human Research Ethics Committee (review reference: 520241317756262). In line with our pre-registration, one participant was excluded due to poor EEG data (>4 excessively noisy electrodes throughout the recording), and four others were excluded because of excessively noisy eye tracking data. Our final sample (*N* = 25) consisted of 21 women and four men aged between 17 and 30 years (*M* = 20.56 years, *SD* = 3.87 years).

### Apparatus

We ran the experiment in a dimmed and electronically shielded room. Stimuli were presented on a HP 27" X27q monitor (2560 × 1440 px resolution), set to a 60 Hz refresh rate, using version 2022.2.5 of PsychoPy (Peirce et al., 2019). Participants used a chin rest at a viewing distance of ∼83 cm and responded via an RB-830 Button Box (Cedrus). We recorded neural data using Curry 7 from 66 EEG channels (64 scalp, two mastoid) connected to a SynAmps RT amplifier (Compumedics Neuroscan) at a sampling rate of 1000 Hz. Channels were laid out according to the 10-10 International Standard montage using an EasyCap (Neuospec AG). AFz and M1 were set to ground and the online reference, respectively. We also recorded left eye movements using an EyeLink 1000 eye tracker (SR Research Ltd.) at a sampling rate of 1000 Hz.

### Multiple Object Monitoring Task

We adapted the MOM task used previously by Karimi-Rouzbahani et al. (2021). In the present version (MOM 2.0, see Figure1A), there were 50 red and 50 blue dots randomly intermingled per block (∼0.5° radius), which onset either from the top left or bottom right of a uniform grey background (∼17.21° onset eccentricity, equal number of red and blue dots per onset location) and smoothly moved along a single visible trajectory on each side towards the centre (∼11.63°/s), which contained a white circular obstacle (∼0.75° radius). Dot onsets pseudorandomly jittered between 900 and 1350 ms to ensure that more than one dot was always present, and only one dot reached the deflection point at a time. At the start of each block, one of the two colours (red or blue) was cued as the task relevant colour (see below) and participants were instructed to fixate on the central obstacle throughout the task.

After 1167 ms post-onset, the dots reached a visible deflection point (∼3.64° eccentricity) where 48% of them were automatically deflected by 90° from their original trajectory, in a random direction, before smoothly offsetting over 833 ms (‘events’). The remaining dots continued their trajectory until either a response was made, or they reached the central obstacle 1500 ms post-onset. If a dot failing to deflect was the relevant colour (a ‘target’), participants had to press a button (333 ms response window). This caused the dots to manually deflect and then smoothly offset over 833 ms. If the dot failing to deflect was the task-irrelevant colour (a ‘distractor’), they had to ignore it. There were an equal number of targets and distractors per block. Dots that reached the central obstacle (i.e., missed targets and correctly rejected distractors) continued along a straight trajectory and smoothly offset over 833 ms.

Auditory feedback was provided following each button press. Correctly responding to target dots before they reached the central obstacle (hits) resulted in a 440 Hz tone. Early responses (before the dot reached the deflection point), late responses (after the dot collided with the obstacle), and false alarms (to either task-relevant or task-irrelevant events) all resulted in a 256 Hz tone. Each button press was attributed to the dot closest to the central obstacle, if it had not yet been responded to, and multiple responses could not be attributed to the same dot.

### Procedure

The experimental session was divided into runs. The first run served as a practice run, where no data were recorded. During each run (both practice and main), participants completed a task block that was not relevant for the present study, and then two MOM task blocks. The ordering of task-relevant colours (red or blue) was randomised such that both served as targets within a given run. Each session lasted two hours, including participant setup and practice. We recorded between seven and ten main runs per participant but only used data from the first seven runs to ensure each participant contributed an equal amount of data to our analyses.

### Identifying Behavioural Outliers

Participants were to be excluded if their overall miss rate deviated from the group mean by more than three standard deviations. No participants met this criterion.

### Behavioural Analysis

Our behavioural analyses first tested whether there were any linear trends in miss rates, false alarm rates to task-relevant dots, or reaction times to successfully deflected targets (hits) as a function of time-on-task. As our EEG-based analyses collapsed across data from all 14 blocks (see EEG Analyses), it was important to check if there were any changes in behaviour over time. Here, false alarms were coded as any response to a task-relevant dot before it reached the deflection point (i.e., early responses), as well as responses to task-relevant *events* (dots that automatically deflected and were therefore not targets).

For each participant, the behavioural measures (miss rate, false alarm rate, and correct reaction times) were averaged within each block. We then separately modelled each measure’s linear relationship with time-on-task (i.e., block number) using univariate linear regression at the participant level. This resulted in participant-level beta weights (slopes) that characterised how much each behavioural measure changed as a function of time-on-task. Group level inference was conducted on these beta weights.

### Preprocessing

We preprocessed the EEG and eye tracking data offline using version 1.6 of MNE-Python (Gramfort et al., 2013) within a Python 3.9.16 environment. First, we integrated the raw eye tracking and EEG data into one data structure, and then concatenated these across runs. Next, the EEG data were re-referenced to the difference between M1 and M2 using the *add_reference_channel* and *set_bipolar_reference* functions. We then removed EEG line noise using a 50 Hz finite impulse response filter and visually identified excessively noisy channels, which were removed and then interpolated using the spherical spline method (Perrin et al., 1989). The data were then epoched from -500 to 1500 ms relative to the onset of each dot. No baseline correction was applied due to the continuous nature of visual stimulation. The epoched data were then downsampled to 250 Hz. No further preprocessing steps were applied.

### EEG Decoding Analyses

#### Decoding Task-Critical Information: Time-to-Deflection Point

We aimed to decode aspects of the MOM task from EEG data that correlated with good performance. Following Karimi-Rouzbahani et al. (2021), we used the time until a given dot reached the deflection point (equivalent to its distance from the deflection point) because this covaried with when dots became increasingly relevant for behaviour.

To do this, we segmented our preprocessed EEG epochs into 15 equally spaced time bins spanning the duration a given dot travelled between onset and deflection point (77.8 ms of data per bin; Figure 1B). We then averaged channel amplitudes over time within each bin, resulting in 63 features per bin (one for each channel). To determine how much dissociable information there was within each time bin, we conducted a series of pairwise, multivariate comparisons between each bin combination (105 combinations total) using five-fold cross-validated linear discriminant analysis at the participant level. Importantly, this was constrained so that data belonging to the same dot across compared bins were kept within the same cross-validation fold. Time bins were also collapsed across onset side (left vs. right), meaning classifiers could not be driven by retinotopic differences between time bins.

Similarly to other studies (Dijkstra et al., 2020; Hetenyi et al., 2024), we quantified the performance of each classifier by the average decision value assigned to each test observation. These values quantified how far away from the decision boundary a given observation was within feature space. We then transformed the values such that they were positive for correctly classified observations and negative for incorrectly classified ones. This metric was chosen over classification accuracy because it allows values further from the decision boundary—where the classifier has more ‘confidence’—to be given more weight when quantifying decoding performance. This process produced a representational dissimilarity matrix (RDM) per epoch characterising the dissimilarity in neural activity between time bins as the associated dot approached the deflection point (Figure 1C). We then further averaged across pairwise comparisons containing a common time bin (e.g., for bin 1, average of: bin 1 vs. 2, bin 1 vs. bin 3, …, bin 1 vs. 15), transforming each RDM into a pseudo-time series from dot onset to when it reached the deflection point (see Figure 1D). The same method was separately applied to three pre-stimulus time bins, giving us a benchmark for effect sizes observed due to chance using this method.

This procedure was applied separately for data evoked from dots that were task-relevant and task-irrelevant under the hypothesis that there would be more neural dissimilarity between time bins for task-relevant dots. Only correctly rejected task-irrelevant dots were included while all task-relevant dots were included. Moreover, we only analysed data before the deflection point (before any responses were required) meaning there were no physical differences in dot stimuli between levels of task-relevance.

### Error Generalisation Analysis

Next, we tested whether the neural dissimilarity measure described above differed between EEG recorded before targets were either hit (i.e., successfully deflected) or missed, using an error generalisation analysis (Karimi-Rouzbahani et al., 2021; Robinson et al., 2022; Woolgar et al., 2019). This method requires very little data on error trials (misses) relative to the training set of correct trials (hits), making it ideal for tasks where errors are infrequent.

Specifically, for each participant, the procedure described above was used to train classifiers to decode *time-to-deflection-point* using a large subset of the hit epochs. We then applied said classifiers to both the miss epochs and an equally-sized, left-out subset of hit epochs in a cross-validated manner. Importantly, there were no physical differences in stimuli before the deflection point, meaning any differences between hits and misses must have reflected internally-driven, state-dependent differences in EEG patterns. We predicted worse decodability for data preceding when participants missed targets compared to (left-out) hit trials, primarily reflecting poorer stimulus encoding during attentional lapses, noting that part of the neural dissimilarity between hits and misses is likely to reflect motor preparation differences.

As in Karimi-Rouzbahani et al. (2021), the number of folds used for cross-validation varied per participant. Specifically, this equalled the number of hit epochs divided by the number of miss epochs. This ensured no differences in the number of test epochs between hits and misses per fold. On average, each participant had 19.76 (*SD* = 9.88) cross-validation folds.

### Predicting Trial-Wise Behaviour from Neural Data

Our main analysis concerned predicting the behavioural outcome to targets on a trial-by-trial basis (i.e., hits vs. misses) from EEG recorded before each dot reached the deflection point. To do this, we employed a hierarchical classification approach that built upon our error generalisation analysis. Specifically, we trained classifiers to decode *time-to-deflection-point* from all but two folds of the hit epoch data (as opposed to just one, like in the above analysis). We then applied these classifiers to the two left-out hit epoch folds (used as validation and test sets, respectively, see below) and the miss epochs. If decodability during this first step was low, the corresponding trial was classified as a miss (otherwise, a hit).

To elaborate, we iteratively included data from each time bin as the dot approached the deflection point, and classified whether it belonged to the correct time bin using the 14 relevant pairwise classifiers (i.e., the true time bin vs. each other time bin). Again, we used distance from the decision boundary as our metric for decoding performance (instead of classification accuracy), which quantified how confidently data were classified per pairwise comparison. These ‘classification scores’ were positive for correctly classified time bins and negative for incorrectly classified bins. This resulted in epoch-level matrices of classification scores, however, unlike in the above analyses, these were not necessarily symmetric because data were iteratively classified per time bin (rather than comparing two time bins from the *same* epoch). These matrices were then transformed into smoothed time series of classification scores via cumulative averaging over the matrix columns at the epoch level.

We first applied these steps to one fold of the left-out hit epochs, the *validation set*, to form a benchmark of expected dissimilarity scores typical of EEG data on *hit* epochs. The steps were then applied to the other (independent) fold of left-out hit epochs, the *test subset*, as well as the *miss* epochs. When the epoch-level classification scores fell below the *validation set* benchmark, these timepoints were hierarchically classified as misses, otherwise they were classified as hits. This entire analysis was cross-validated such that each hit epoch fold served independently within the *validation* and *test* sets. The analysis was repeated ten times to obtain a reliable estimate of classification accuracy. The data reported here are the average across repeats.

#### Optimising the Decision Boundary

We then aimed to further optimise this form of hierarchical classification using the same approach as Karimi-Rouzbahani et al. (2021). Importantly, this was completed in a leave-one-participant-out manner to avoid circularity when assessing its efficacy. For all but the left-out participant, we completed the approach detailed above, however, this time, the cumulatively averaged validation set time series (i.e., the benchmark) was lowered by its standard deviation across trials, scaled between 0.1 and 4 (steps of 0.1). Specifically, a scaler was chosen such that hits and misses were best separated per participant. The scaler that performed best (on average) within the training participant data was then applied to the left-out participant.

#### Pre-registration Deviation

Please note, the hierarchical classification method described here slightly deviates from what we originally pre-registered. Specifically, the data from each time bin were only going to be compared against data from all future time bins when constructing the trial-level accumulated classification score time series (see Figure 6 in Karimi-Rouzbahani et al., 2021). As can be seen within Supplementary Figure 1, this approach proved ineffective at dissociating hits and misses. We then iteratively compared the data from each time bin to *all* other time bins (i.e., the immediate past and future), increasing the number of comparisons made per epoch. Indeed, this change proved effective at dissociating hits and misses (see Results).

### Statistical Inference

All hypotheses were tested in version R2024a of MATLAB (MathWorks) using paired samples Bayesian *t*-tests (Rouder et al., 2009). These quantified the amount of evidence in favour of the alternative hypothesis relative to the null hypothesis in the form of a Bayes factor (BF_10_). All tests employed a Cauchy prior (half-Cauchy for one-tailed tests) with default width (*r* = 0.707). As per the recommendations of Teichmann et al. (2022), we included a null interval for tests involving neural data. For two-tailed tests, the null interval ranged between δ = -0.5 and δ = 0.5 (Moerel et al., 2022). For one-tailed tests, it ranged between δ = 0 and δ = 0.5 (Lowe et al., 2023). BF_10_ values falling between 1/3 and 3 were deemed to have inconclusive evidence for either the null or alternative hypotheses.

### Eye Tracking Analyses

Decoding results can be driven by systematic eye movements (Quax et al., 2019). While participants were instructed to fixate on the central obstacle, we nonetheless checked that our results could not be driven by eye movements. Specifically, we repeated the above three analyses using the gaze position and pupil size data instead of the EEG data. These found support for the null (Bayesian statistics) for all effects (see Supplementary Material). Thus, we are confident that our EEG-based decoding results were driven by neural activity related to the task and not eye movement-related artefacts.

### Exploratory Analysis: Modulations in Posterior Alpha Power

Finally, we ran an exploratory, frequency spectra-based analysis. As highlighted in the Introduction, previous work, using static stimuli presented in isolation, has shown that trial-wise increases in posterior alpha power correlate with poor task performance in sustained attention tasks (Mazaheri et al., 2009; O’Connell et al., 2009). More generally, posterior alpha power has been shown to decrease within the hemisphere contralateral to the locus of spatial attention (Foxe & Snyder, 2011; Ikkai et al., 2016; Sauseng et al., 2005), which is largely thought to reflect the disinhibition of task-relevant stimulus encoding. Thus, we tested whether this univariate measure also correlates with different aspects of our MOM task (i.e., whether a dot was task-relevant and whether targets would be detected).

Each dot-specific EEG epoch was re-referenced to the average activity across all channels and then baseline corrected using the data evoked while dots travelled along the trajectory before reaching the deflection point (i.e., 0 to 1167 ms post stimulus onset). We then randomly subsampled the epochs such that trial numbers were equal between levels of task-relevance (task-relevant vs. task-irrelevant dots) and outcome (hit vs. miss) within each side of the display (left and right onset, respectively). We ran a Fast Fourier Transform (SciPy, version 1.12) on the data recorded while each dot was within the second-half of its trajectory before reaching the deflection point (i.e., from 548 ms before the deflection point; 2 Hz resolution). We then modelled the isolated periodic activity between 3 and 55 Hz across individual channels per epoch using version 1.1 of the ‘*fitting oscillations & one over f*’ (*FOOOF*) algorithm (Donoghue et al., 2020). Here, the peak width limits were set between 4 and 12 Hz, and the aperiodic mode was fixed. Because we expected to observe effects within parietal and occipital electrodes, we averaged the modelled alpha-band power (8 to 12 Hz) across epochs within a left (P1, P3, P5, P7, PO3, PO7, and O1) and right (P2, P4, P6, P8, PO4, PO8, and O2) region of interest (ROI), and entered these values into our statistical models.

### Analyses of Variance

We ran two sets of 2x2 Bayesian repeated measures ANOVAs (Rouder et al., 2012) using version 0.18.3 of JASP (JASP Team). Briefly, these quantify whether including each factor, and the interaction term, increases the amount of variance explained within a general linear model in the form of a Bayes factor (BF_inc_). The first ANOVA included task-relevance (task-relevant vs. task-irrelevant) and ROI relative to dot onset (ipsilateral vs. contralateral) as factors. We only included data before dots were either hit or correctly rejected in this analysis. The second ANOVA was restricted to data evoked before target dots reached the deflection point. This included behavioural outcome (hit vs. miss) and hemisphere (ipsilateral vs. contralateral) as factors. Interactions were followed up using two-tailed Bayesian paired samples *t*-tests with a Cauchy prior (*r* = 0.707).

## Results

Across blocks, participants successfully deflected (hit) an average of 93.64% of targets (*SD* = 3%; missing 6.66% of targets), with a mean hit reaction time of 229.69 ms (*SD* = 29.77 ms) relative to when the dot passed the deflection point. They made false alarm responses to 22.28% (*SD* = 11.34%) of task-relevant events and 0.08% (*SD* = 0.25%) of task-irrelevant events (when, in both cases, the dot deflected automatically). They responded to very few distractors (dots that did not automatically deflect but were the task-irrelevant colour; *M* = 0.21%, *SD* = 0.39%). Finally, participants made very few ‘too early’ responses (before the dot reached the deflection point) to either task-relevant (*M* = 0.85%, *SD* = 1.75%) or task-irrelevant dots (*M* = 0.6%, *SD* = 1.76%). All remaining dots were correct rejections of events and distractors (i.e., participants correctly did not press the button).

Figure 2 shows how miss rates, false alarm rates to task-relevant dots, and reaction times to successfully deflected targets (i.e., hits) linearly changed over the course of the experiment. There was inconclusive evidence for whether or not miss rates (BF_10_ = 1.64) and reaction times (BF_10_ = 0.5, direction of the null) changed over time. There was no effect of time on task (null hypothesis support) on false alarm rates to task-relevant dots (BF_10_ = 0.32).

**Figure 2.**
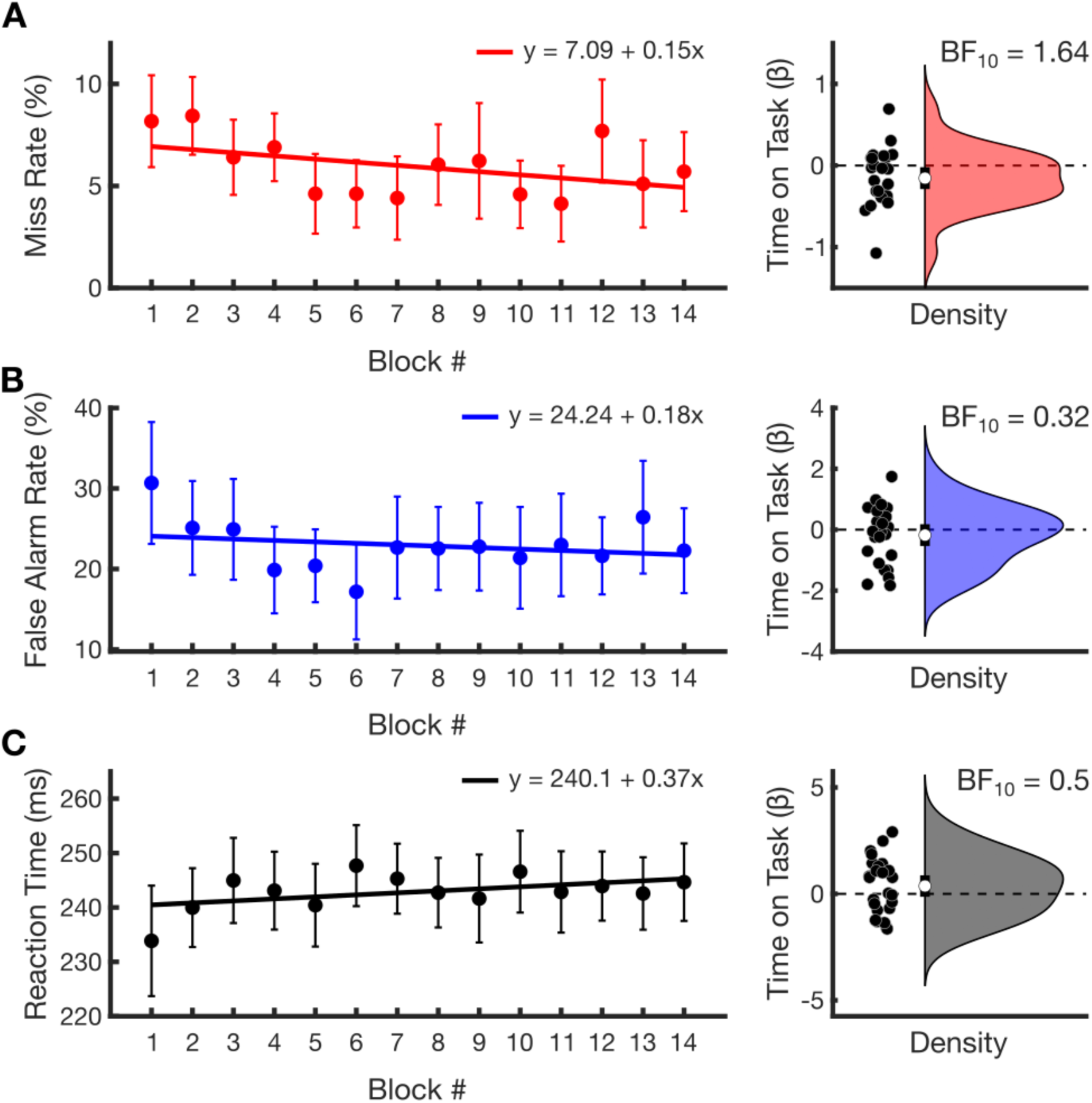
Behavioural trends across task blocks. **A)** The left subplot depicts group-averaged miss rates over the course of the experiment. Error bars denote 95% confidence intervals (*N* = 25). The fitted line is made from the average participant-level parameter estimates derived using univariate linear regression. Within the line equation, x refers to ‘block #’ and y refers to ‘miss rate (%)’. The right subplot depicts the distribution of participant-level slopes (beta weights), characterising how miss rates linearly changed over the course of the experiment for each participant. Points represent individual participants, and the horizontal dotted line marks a slope of zero (i.e., no change over time). The average beta weight and 95% confidence interval is depicted by the white circle and vertical line, respectively. The Bayes factor (BF_10_) testing whether these slopes differed from zero is shown next to each distribution. **B)** and **C)** depict false alarm rates to task-relevant dots (%) and correct reaction time (ms), respectively, using the same layout as **A**.

### Decoding Time-to-Deflection-Point by Task-Relevance

First, we tested whether task-relevant dots evoked more *time-to-deflection-point* information (i.e., greater dissimilarity between time bins) compared to task-irrelevant dots—as should happen if the distinction between time bins reflects neural coding of an attended stimulus. This was evident across almost all time bins (Figure 3A-C). Surprisingly, we found the opposite effect during the first time bin post-onset, which we suspect reflects inhibition of task-irrelevant dots (see Discussion).

**Figure 3.**
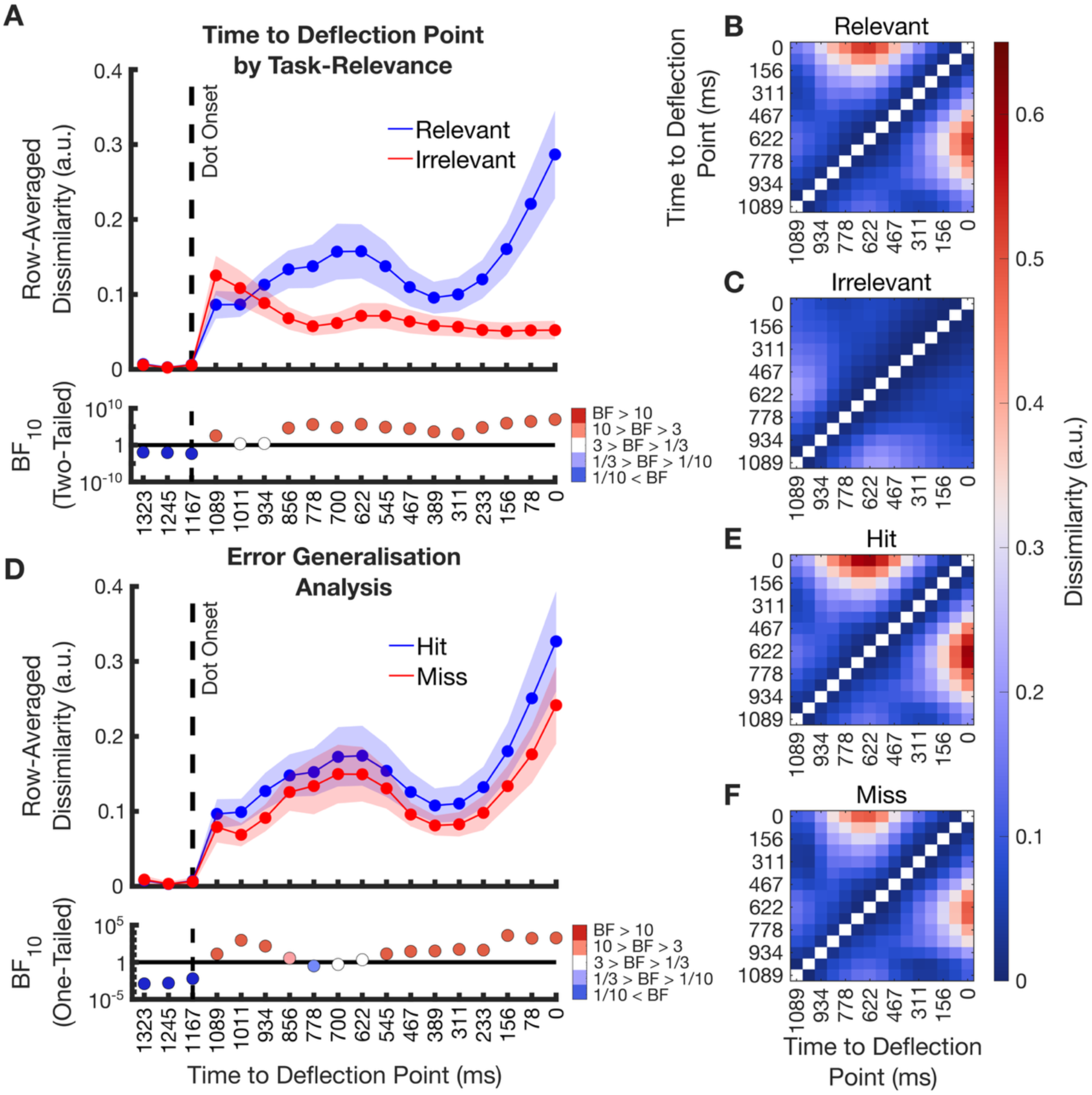
Neural dissimilarity between time bins. **A)** The pseudo time-series quantifying the cross-validated average dissimilarity in EEG data between time bins evoked by task-relevant (blue line) and task-irrelevant (red line) dots as they approached the deflection point. Shading denotes 95% confidence intervals (*N* = 25). Dot onset is marked by a dashed vertical line. Time points before this line were made using three ∼78 ms time bins of EEG data to provide a benchmark of effect sizes that can be observed due to chance using this method. Bayes factors (BF_10_) quantify whether task-relevant and task-irrelevant dots evoke different amounts of dissimilarity. The horizontal line marks a BF_10_ of 1, filled circles show BF_10_ with sufficient evidence for interpretation. **B-C)** Representational dissimilarity matrices (RDMs) that quantify the pairwise dissimilarity between multivariate patterns of EEG evoked whilst the dots were within each time bin. Warmer colours index greater dissimilarity. The diagonal is left empty as EEG data were not compared within time bins. These RDMs were used to construct the data presented in **A**. **D**-**F)** show the error data analysis results wherein classifiers were trained on hit data and tested on either miss data (red line) or an equally sized portion of left-out hit data (blue line) with the same conventions as for **A-C**.

For task-relevant dots, there was an increase in dissimilarity from ∼311 ms before the deflection point that did not occur for task-irrelevant dots. This seems likely to reflect the increased importance of paying attention to the dot at this critical point along the trajectory as well as possible motor preparation (although note that participants did not yet know whether a response was required; see Discussion). As can be seen within the accompanying RDM (Figure 3B), this increase in dissimilarity was primarily driven by comparisons between the time bins ending 700 and 0 ms before dots reached the deflection point, respectively. It is possible that dissimilarity scores were greatest here because the interstimulus interval between dot onsets (900 to 1350 ms) meant that these two time bins shared no visual overlap, thereby increasing their dissimilarity. However, as this only occurs for task-relevant dots, it cannot be solely driven by retinotopic differences between time bins (retinotopic information is equivalent between task-relevant and task-irrelevant dots). Thus, these data collectively show that during the MOM task, patterns of neural activity representing the critical information about the task (time until deflection) were clearly modulated by whether or not the dot was task-relevant, consistent with effects of attention.

### Error Generalisation Analysis

Next, we tested whether the multivariate patterns of EEG data before misses systematically differed from the data recorded before hits. Because there were so few miss epochs per participant (see Behaviour), directly decoding these data using the typical cross-validation approach is not possible. Instead, we used our error generalisation analysis technique (Karimi-Rouzbahani et al., 2021; Robinson et al., 2022; Woolgar et al., 2019), wherein we trained a series of pairwise classifiers to decode *time-to-deflection-point* during hit epochs and then generalised their decision rules to miss epochs. We expected to see a drop in distance dissimilarity for cross-decoded miss epochs compared to cross-validated hit epochs under the hypothesis that EEG evoked before the deflection point systematically differed between hits and misses.

When comparing the two time series, Bayes factors favoured evidence for the alternative hypothesis across almost all time bins, including the moments immediately following dot onset (Figure 3D). Like above, this difference was greatest immediately before dots reached the deflection point, which is when they were most behaviourally relevant. These data suggest that patterns of EEG preceding the deflection point systematically differed between hits and misses.

### Predicting Trial-Wise Behaviour from Neural Data

As there was dissociable neural activity before hits and misses (Figure 3D), our final planned analysis tested whether we could use this difference to predict behaviour on a trial-by-trial basis before targets reached the deflection point. As a brief reminder, classifiers were first trained on hit data to discriminate between EEG evoked during each time bin prior to when dots reached the deflection point. These classifiers were then tested on left-out subsets of EEG evoked before targets were either hit or missed, resulting in cumulative time series data (Figure 4A-B). As can be seen from Figure 4C, cumulative classification scores were qualitatively greater for hits compared to misses from ∼700 ms before they reached the deflection point, which is well before the participant knew the dot was a target.

**Figure 4.**
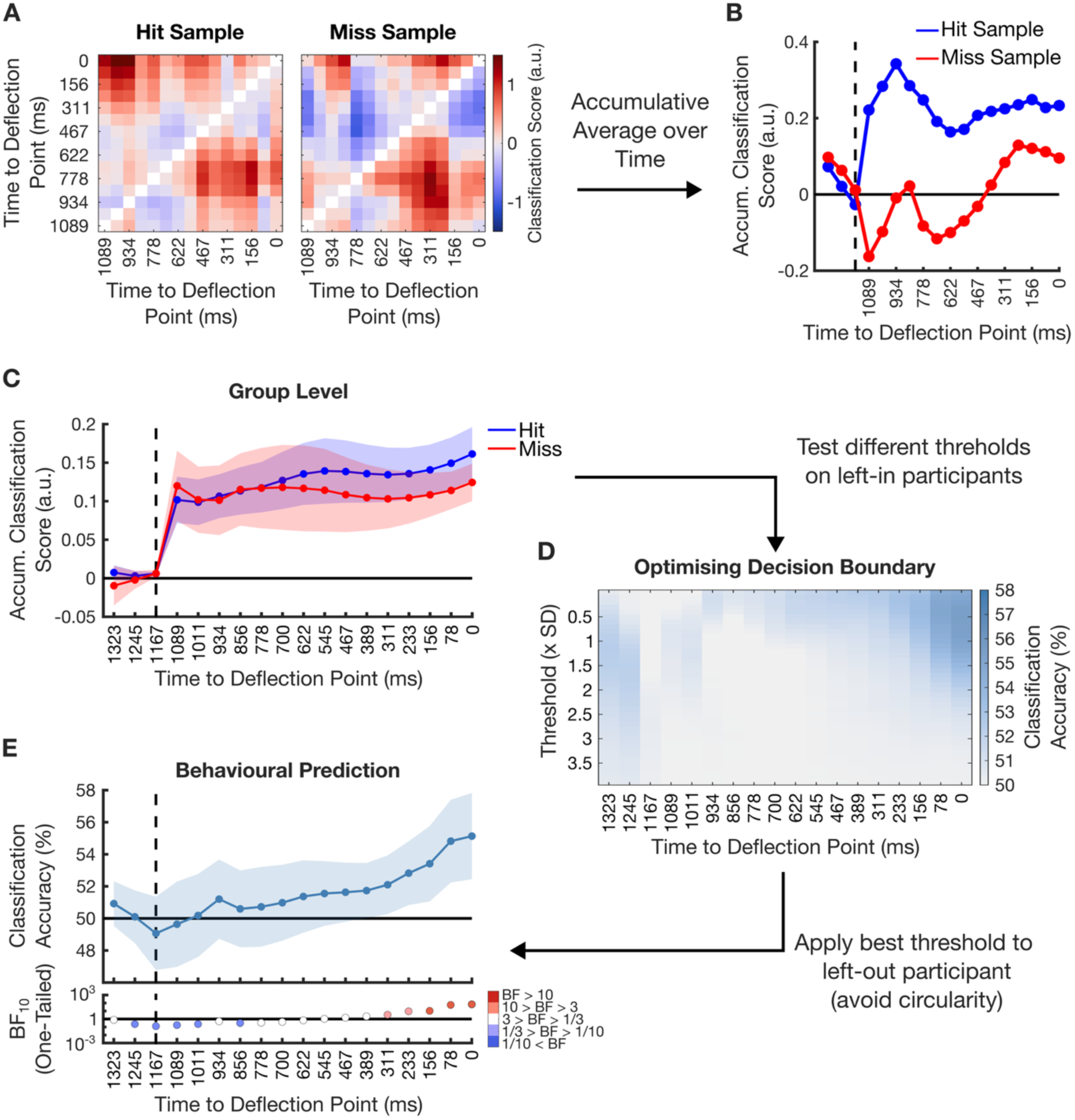
Predicting behaviour from neural patterns. **A)** Single-trial dot data are iteratively classified as being drawn from the correct time bin vs. each other time bin as the dot approaches the deflection point. Classifiers were trained on hit data. Warmer colours denote comparisons where time bins were correctly classified. The (empty) diagonal in each matrix corresponds to the dot’s true time bin. Values are cumulatively averaged across over columns forming the time series within **B**. **C)** Grand-averaged time series using this method. The dashed vertical line denotes dot onset, and the solid horizontal line denotes chance scores. Values are increasingly higher for hits compared to misses from ∼700ms before deflection. **D)** Classification accuracy for predicting behavioural outcome (hit vs. miss) as a function of time to deflection point using our hierarchical classifier when it is set to different threshold values (chance = 50%). **E)** Classification accuracy over time for predicting behaviour as a given dot approaches the deflection point derived using a leave-one-participant-out approach. We iteratively identified an optimal threshold value using the left-in participants and applied it to the left-out participant. Vertical and horizontal lines denote dot onset and chance values, respectively. Shading shows 95% confidence intervals. Bayes factor convention is identical to Figure 2.

On a fold-by-fold basis, we defined a benchmark of cumulative classification scores characteristic of when a target dot was about to be successfully detected from a portion of left out hit epochs (the *validation set*). Target dots evoking cumulative classification scores falling below this benchmark minus a specific threshold (based on the standard deviation within the validation set) were classified as misses, otherwise hits. A matrix of classification accuracy for hierarchically predicting whether a target dot was about to be hit or missed, as a function of threshold values and time bins, is shown in Figure 4D. Classification accuracy, averaged across time bins, was best when the scaler threshold was set to 0.92 (*SD* = 1.34).

Next, we tested whether behavioural prediction classification accuracies were meaningfully above chance before dots reached the deflection points (i.e., whether we could predict when participants were about to make an error). As detailed within the Methods section (see Optimising the Decision Boundary), we used a leave-one-participant-out method, where the best performing threshold across participants was applied to the left-out participant. Classification performance (hit vs. miss) was reliably above chance (BF_10_ > 3) from ∼311 ms before the deflection point (Figure 4E). Our classification accuracy peaked as dots reached the deflection point. Thus, while not as strong as with MEG (Karimi-Rouzbahani et al., 2021), these results demonstrate that the drop in task-based information reported above (see Error Generalisation Analysis; Figure 3D-F) is indeed predictive of when participants were about to make errors within the MOM task on a trial-by-trial basis.

### Exploratory Frequency Spectra Analyses

Our final, exploratory, analysis tested whether posterior alpha power, recorded before dots reached the deflection point, covaried with the important aspects of the MOM task used in the decoding analyses (Figure 5).

**Figure 5.**
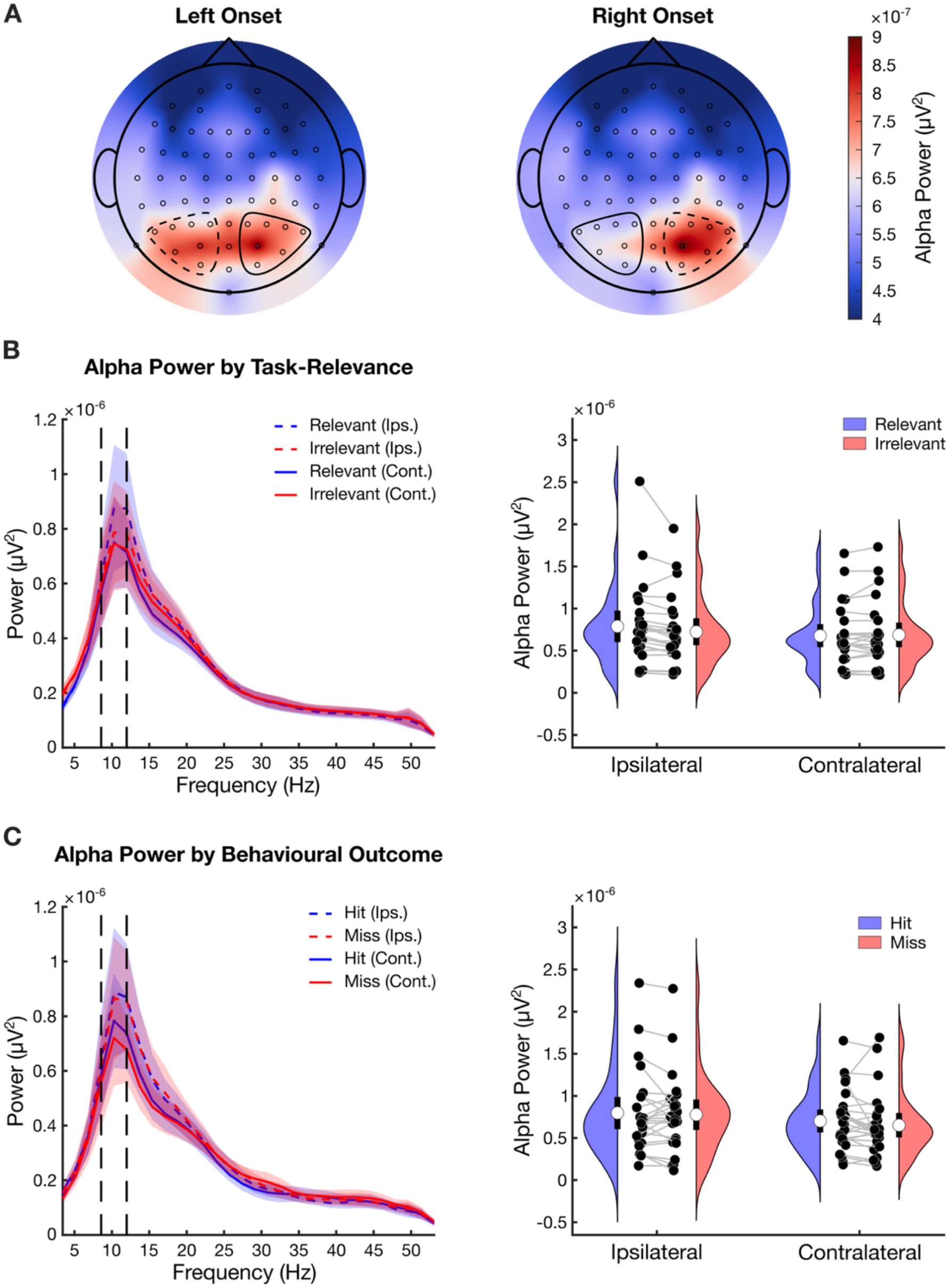
Task-evoked posterior alpha power. **A)** Whole-scalp topographies of periodic alpha power (8 to 12 Hz) evoked as dots approached the deflection point (collapsed across levels of task-relevance). Individual circles represent EEG channel locations. The dashed-lined clusters correspond to the ipsilateral region of interest (ROI). The solid-line clusters correspond to the contralateral ROI. **B)** The left subplot shows posterior periodic spectra split by dot task-relevance and ROI. All four conditions exhibit a clear peak within the alpha frequency band (indicated by vertical dashed lines). Shading shows 95% confidence intervals. Ips. = ipsilateral ROI; Cont. = contralateral ROI. The right subplot shows group-level distributions of posterior periodic power averaged across the alpha band per condition. **C)** Periodic power data evoked before target dots were either hit or missed using the same convention as **B**.

### Alpha Power by Task-Relevance

We found evidence for a main effect of both task-relevance (BF_inc_ = 4.8) and ROI laterality (BF_inc_ = 54.49) on alpha power, modulated by an interaction (BF_inc_ = 15.43). Specifically, task-*relevant* dots evoked more alpha power than task-*irrelevant* dots within the *ipsilateral* ROI (BF_10_ = 3.4), but there was no effect of task-relevance within the *contralateral* hemisphere (BF_10_ = 0.26). Both task-*relevant* and task*-irrelevant* dots evoked more alpha power within the *ipsilateral* ROI compared to the *contralateral* ROI (BF_10_ = 11.56 and BF_10_ = 17.07, respectively).

### Alpha Power by Behavioural Outcome

Regardless of the response, dots that subsequently became targets evoked more alpha within the ipsilateral hemisphere compared to the contralateral hemisphere (BF_inc_ = 6.13). In contrast, we found no difference between hits and misses in the alpha band (BF_inc_ = 0.38), nor was there an interaction (BF_inc_ = 0.39). This suggests that posterior alpha power did not reliably change before targets were hit or missed.

## Discussion

The present study aimed to used neural decoding techniques to objectively identify spontaneous lapses in attention from EEG activity. Specifically, we further developed a recent hierarchical classification method (Karimi-Rouzbahani et al., 2021) to predict when participants would fail to detect (miss) moving target stimuli within a dynamic sustained attention task. Importantly, we only used data from *before* the stimuli required a behavioural response (i.e., before the stimuli reached the deflection point; Figure 1), meaning the information that differentiated between when participants *were about to* miss a target must have been internally driven. We use this change in neural signal as an index of lapses in sustained attention.

Participants missed only a small proportion of the targets, and there were no striking time-on-task effects. This contrasts with situations where targets are rare, whereby performance tends to drop dramatically over time (i.e., ‘vigilance decrements’; see Thomson, Besner et al. 2015). False alarm rates to task-relevant events (responses to cued dots that already deflected) remained stable throughout the experiment. Consistent with previous research, these behavioural data suggest that when targets are frequent, performance can be sustained relatively effectively, with error rates (particularly misses) being low and relatively unaffected by time on task (Karimi-Rouzbahani et al., 2021; Rich et al., 2008; Wolfe et al., 2007). Importantly, because misses were made throughout the experiment (albeit at a low rate), our ability to predict when they were about to occur cannot have been driven by state-based changes correlated with time on task. Instead, the predictions rely on trial-by-trial changes in neural patterns between hits and misses, which we suggest reflects the consequences of a lapse in sustained attention.

Turning now to the neural results, the information we decoded to index sustained attention reflected how long until a given moving dot reached the deflection point, where it might require a response. This ‘time until deflection’ therefore reflects the increasing relevance of the dot for behaviour (as well as its decreasing distance to the deflection point). As a first pass, we showed that there was more *time-to-deflection-point* information for dots cued as task-relevant compared to dots that were task-irrelevant, particularly just before they reached the deflection point. This pattern of results converges well with previous literature showing that (static) stimuli that are attended to or behaviourally relevant are better decoded than those that are not relevant (Goddard et al., 2022; Jehee et al., 2011; Jiang et al., 2013; Moerel et al., 2022; Smout et al., 2019; Woolgar et al., 2015).

Surprisingly, there was more *time-to-deflection-point* information for task-irrelevant dots than task-relevant dots during the earliest time bins (Figure 3A). Because dot onset is highly salient (Van Pelt et al., 2025), and some initial processing is required before a dot can be deemed irrelevant and then ignored, it seems likely this effect reflects distractor suppression processes (Gaspelin & Luck, 2018). As our analysis depended on the average dissimilarity of a given time bin compared with all the other time bins (see Figure 1B-D), it seems likely that there would be a difference for task-irrelevant dots between the initial time bins, during which attention would have been drawn to the dot, from subsequent time bins, where there is little reason to continue tracking it. By contrast, participants were required to sustain their attentional resources on task-relevant dots throughout, effectively making the earlier and later time bins less dissimilar than for irrelevant coloured dots.

Most important for our research question, the decoding *decreased* when classifiers trained on data from when targets were *about to be* detected (i.e., hit) were tested on data preceding when targets were missed (Figure 3D-F). This result can only occur if the neural patterns evoked before participants detected targets were reliably weaker or absent when targets were missed. While our error generalisation analysis cannot directly test whether missed dots are represented more poorly than hit ones or just represented differently, previous studies have shown that visual cortex is less responsive, or increasingly inhibited, when (static) target stimuli are not detected (Cohen & Maunsell, 2010; Ergenoglu et al., 2004; Mazaheri et al., 2009; O’Connell et al., 2009; Thut et al., 2006; van Dijk et al., 2008; Weissman et al., 2006). Our findings demonstrate that less of the critical task-information is decodable when participants go on to miss a target, compared to when they successfully respond, consistent with a decrease in neural encoding.

Based on the difference between hits and misses in neural patterns, we then tested whether we could predict misses before they occurred from the *time-to-deflection-point* information. Our second level classifier could reliably (above chance) determine whether a miss was about to occur from ∼311 ms before dots reached the deflection point (Figure 4E). Crucially, this was at the single-trial level, which is a necessary level of specificity if we are to develop objective measures of sustained attention and be able to detect transitory lapses. Our classification accuracy was lower than that achieved with MEG data (Karimi-Rouzbahani et al., 2021), but this is unsurprising given the difference in sensitivity between EEG and MEG (see Fred et al., 2022). Regardless, these data provide proof-of-concept that an objective measure of sustained attention can be extracted from ongoing EEG activity, serving as a foundation for future development.

When participants were successfully tracking a task-relevant dot, it seems likely that they would be engaging in anticipatory motor preparation, in case it was a target. Within the MOM task, target detection requires not only perceptual processing but also the readiness to execute a response, due to the tight timeframe to prevent a collision (333 ms). Such motor preparation would increasingly influence neural activity as the dot approached the deflection point. On trials when participants failed to track a task-relevant dot, defined by missing the target, this motor preparation was presumably less or absent. Thus, it seems likely that differential motor preparation contributes to the difference in dissimilarity in neural data between time bins for hits versus misses. Such differential motor preparation contributing to the prediction of hits and misses is not a confound because our goal was to identify neural signatures that correlate with misses, so that they can serve as an objective index of attentional lapse on a trial-by-trial basis. A change in readiness to respond is a logical consequence of a lapse in attention as the dot approached the deflection point.

In addition to our main pre-registered analyses, we explored whether posterior alpha power covaried with aspects of the task and subsequent behaviour (Figure 5). Oscillatory alpha rhythms have been postulated to have a functional role in suppressing unattended regions within the visual field (Foxe & Snyder, 2011; Ikkai et al., 2016; Sauseng et al., 2005; however, see Foster & Awh, 2019). Moreover, there is reason to suspect that poor stimulus encoding and target detection (i.e., consequences of attentional lapses) might correlate with spontaneous increases in posterior alpha power (Griffiths et al., 2019; Mazaheri et al., 2009; O’Connell et al., 2009; Thut et al., 2006; van Dijk et al., 2008).

In alignment with previous work, we found more alpha power within the hemifield ipsilateral to a given dot’s location on the screen, which, according to dominant theory (Foxe & Snyder, 2011; Ikkai et al., 2016; Jensen, 2024; Jensen & Mazaheri, 2010; Sauseng et al., 2005), reflects suppressed neural activity within the visual field that is less relevant for behaviour (here the side without a dot of the relevant colour approaching the deflection point). Moreover, this hemispheric difference was greater for task-relevant dots, suggesting participants were more spatially selective for these compared to task-irrelevant dots. This aligns with our decoding analysis showing that task-relevant dots evoked more *time-to-deflection-point* information than task-irrelevant dots (Figure 3A-C). Despite this, we found no differences in alpha power between task-relevant and task-irrelevant dots within hemispheres contralateral to the dot onset. If we accept that the increases in alpha found here did indeed reflect distractor suppression, this suggests that participants always selected the dot closest to the deflection point, regardless of whether it was task-relevant or not, but the degree to which the opposite side of the display was suppressed depended upon whether the dot colour was currently relevant.

Finally, despite finding clear effects of task-relevance, we found no differences in posterior alpha power during the moments before a dot was either hit or missed (Figure 5C). This contrasts with our decoding approach, which found differences in *time-to-deflection-point* information between outcomes (Figure 3D-F). As such, our error generalisation and hierarchical classification analyses could provide more effective tools for identifying misses over than the more classical approach of analysing frequency spectra.

## Conclusion

The overall goal of this study was to objectively index lapses in sustained attention from EEG data whilst participants monitored a moving visual display. To this end, we further developed a hierarchical classification algorithm that exploits decreases in task-critical information to predict when participants are about to miss targets on a trial-by-trial basis. This approach provided small but reliable predictions of behaviour based on the neural pattern of activity. We did not find any meaningful differences for classical indices of attention lapses (here, increases in posterior alpha power). Understanding what changes when attention is successfully maintained versus lapsing is a crucial area of research and may inform real-world applications to reduce fatal errors. These results serve as a foundation for the further development of sensitive and specific methods to objectively detect lapses in sustained attention based on patterns of brain activity.

## Supplementary Material

### Pre-registration Deviations

As mentioned within the main text, this study was pre-registered on the Open Science Framework (https://osf.io/cu87z). We made some minor deviations from our pre-registered analyses. The rationale for each change is detailed here.

### Task Changes

Due to a coding error, 52% of task-relevant dots and 52% of task-irrelevant dots were targets and distractors, respectively, instead of 50%. However, as these were still matched between relevant and irrelevant dot colours, this did not affect the integrity of the study or the results.

### Behavioural Analyses

We planned to use perceptual sensitivity (*d*’) as our measure for task performance. However, this metric was difficult to calculate due to there being multiple sources of false alarms: responding to event dots after they had automatically deflected (regardless of task-relevance), responding to any dot before it reached the deflection point, and responding to distractors (see Figure 1). Thus, sensitivity slightly varied depending on which false alarm type(s) were included or omitted during its calculation. Because we did not pre-register exactly how sensitivity would be derived, and this degree of experimenter freedom could have potentially changed our results, we opted for the more simplistic measure of miss rates and false alarms to be reported separately.

### Stopping Rule

We initially planned to check the results after 20 participants to see if we had sufficient data. However, as this experiment forms a control group for another experiment examining the effects of low target frequency (https://osf.io/wxfkd), we ended up collecting 30 participants before analysing the data. Of these, 5 had to be excluded based on our pre-registered exclusion criterion, leaving us with 25 participants.

### Dissimilarity as a Metric for Decoding Performance

We pre-registered that decoding *time-to-deflection-point* information would be quantified using classification accuracy. A more recent approach, however, suggested that using the average decision value assigned to each observation is a more fine-grained estimate of decoding performance (Hetenyi et al., 2024), so we used this slight modification instead.

### Results from Pre-registered Behavioural Prediction Approach

As explicitly detailed within the main text (see Methods), we slightly deviated from our pre-registration when predicting trial-wise behaviour. As a brief reminder, the results within Figure 4 of the main text were acquired by iteratively comparing data from each time bin to *all* other time bins. Within our pre-registration, we stated that data would only be compared to *future* time bins, however, we were unable to predict behaviour using this approach (see Supplementary Figure 1).

**Supplementary Figure 1.**
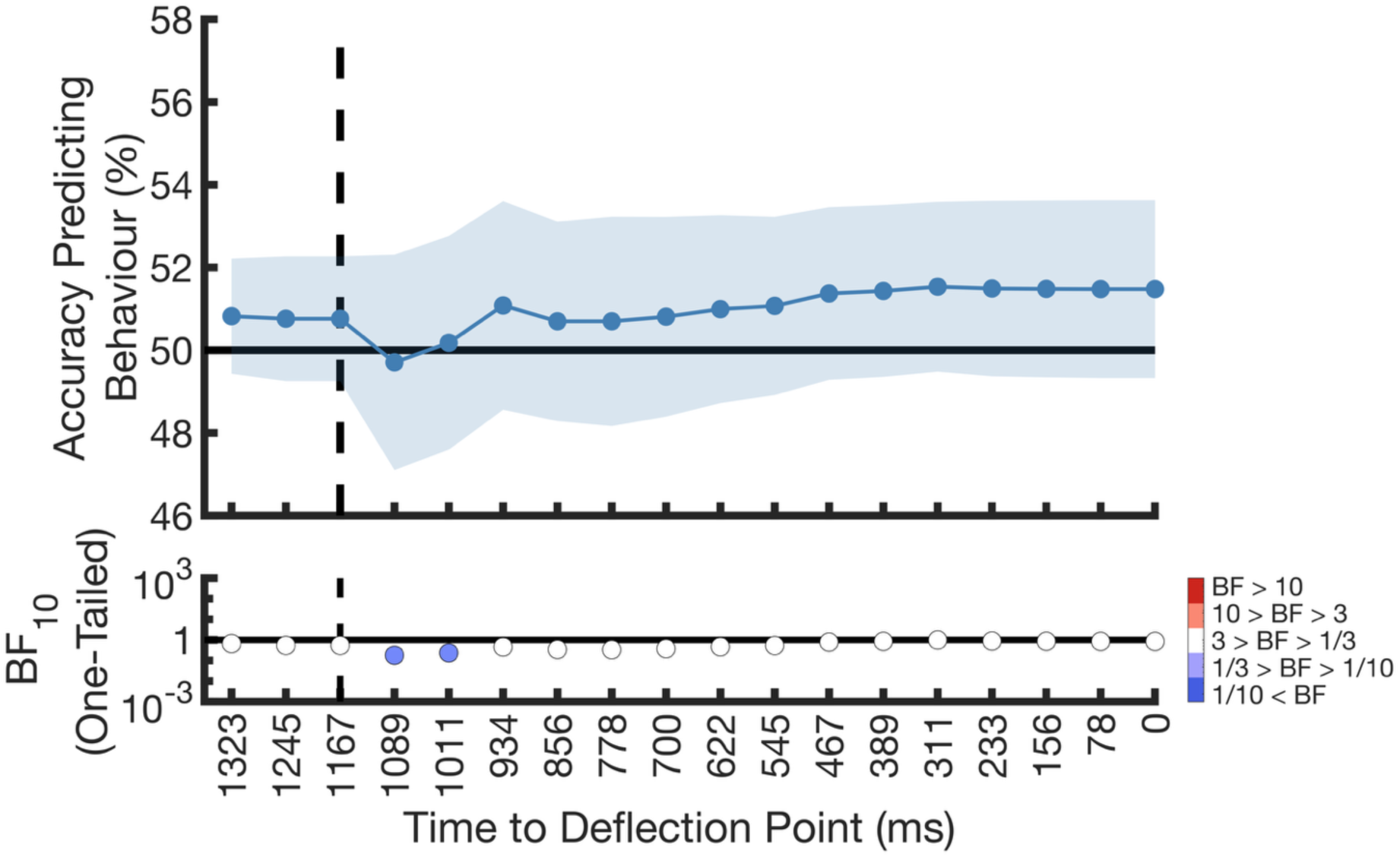
Null results for our pre-registered behavioural prediction approach. Classification accuracy over time for predicting behaviour as a given dot approaches the deflection point using our originally pre-registered approach. This was derived using a leave-one-participant-out approach, where we iteratively identified an optimal threshold value using the left-in participants and applied it to the left-out participant, as we did for the EEG data. The dashed vertical and solid horizontal lines denote dot onset and chance values, respectively. Shading shows 95% confidence intervals (*N* = 25). Bayes factors (BF_10_) quantify the evidence for classification accuracy above chance. The horizontal line marks a BF_10_ of 1, filled circles show BF_10_ with sufficient evidence for interpretation (in this case showing evidence for the null).

**Supplementary Figure 2.**
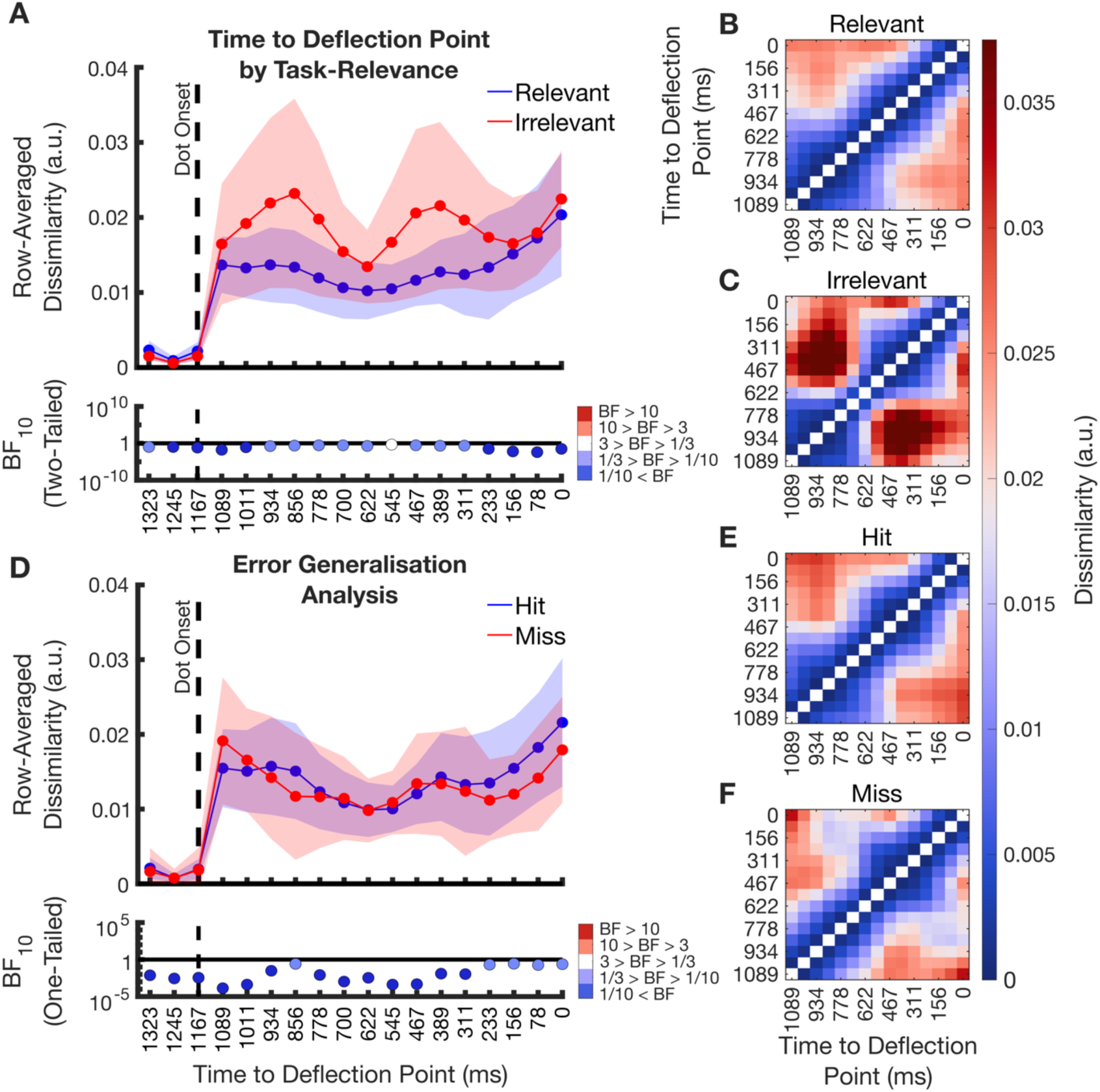
Eye tracker dissimilarity between time bins. **A)** The pseudo time-series quantifying the cross-validated, average dissimilarity in eye tracking data (gaze position and pupil size) between time bins evoked by task-relevant (blue line) and task-irrelevant (red line) dots as they approached the deflection point. Shading denotes 95% confidence intervals (*N* = 25). Dot onset is marked by a dashed vertical line. Time points before this line were made using three ∼78 ms time bins of data to provide a benchmark of effect sizes that can be observed due to chance using this method. Bayes factors (BF_10_) quantify whether task-relevant and task-irrelevant dots evoke different amounts of dissimilarity. The horizontal line marks a BF_10_ of 1, filled circles show BF_10_ with sufficient evidence for interpretation. **B)** and **C)** show the representational dissimilarity matrices (RDMs) that quantify the pairwise dissimilarity between multivariate patterns of eye tracking data whilst the dots were within each time bin. Warmer colours index greater dissimilarity. The diagonal is left empty as data were not compared within time bins. These RDMs were used to construct the data presented in **A**. **D)**, **E)**, and **F)** show the error data analysis results wherein classifiers were trained on hit data and tested on either miss data (red line) or an equally sized portion of left-out hit data (blue line) with the same conventions as for **A**. Note that, unlike EEG, Bayes factors found no differences between conditions from eye tracking data.

**Supplementary Figure 3.**
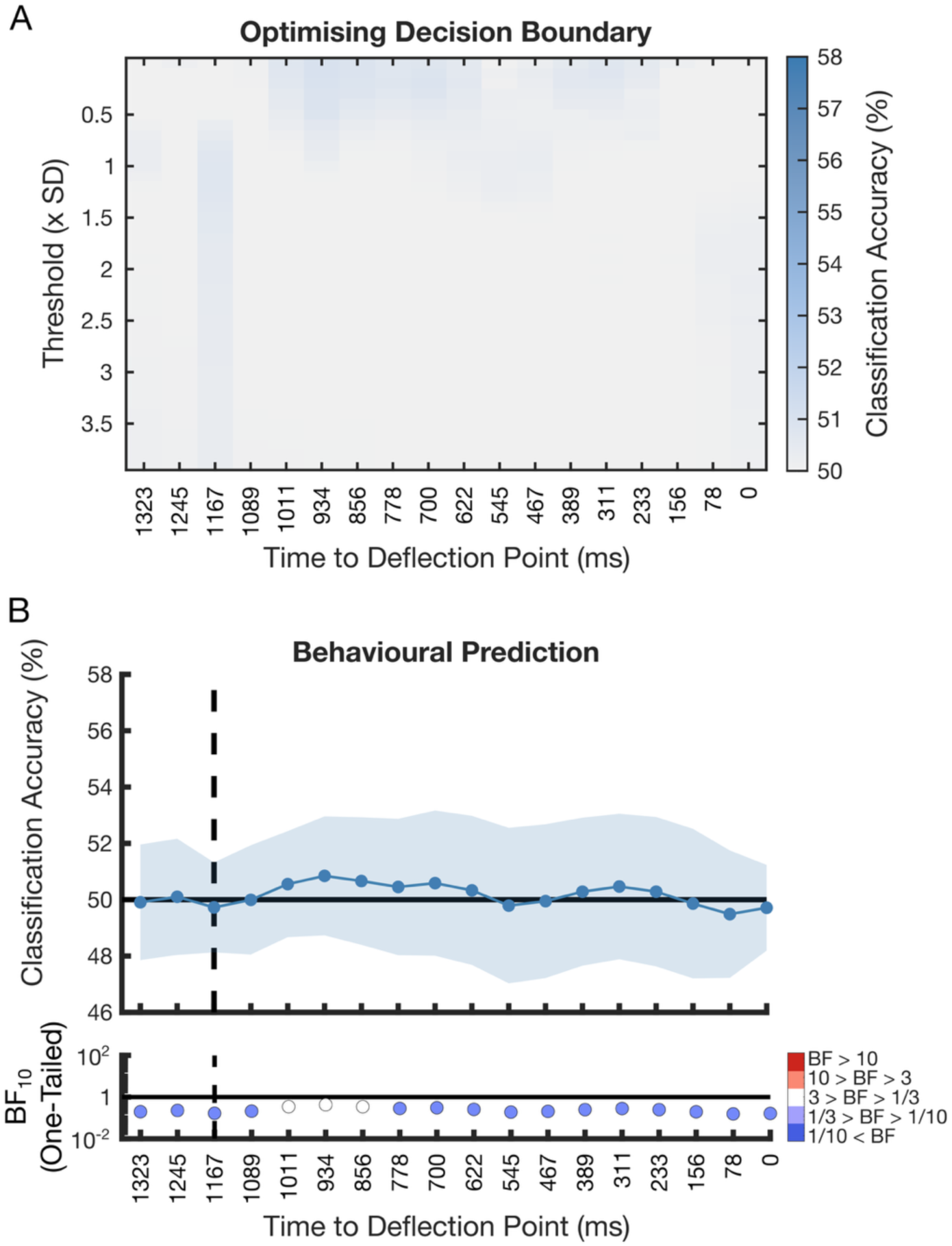
Behavioural prediction results from eye tracking data. **A)** Classification accuracy for predicting behavioural outcome (hit vs. miss) as a function of time to deflection point using our hierarchical classifier when it is set to different threshold values (chance = 50%). **B)** Classification accuracy over time for predicting behaviour as a given dot approaches the deflection point. This was derived using a leave-one-participant-out approach, where we iteratively identified an optimal threshold value using the left-in participants and applied it to the left-out participant, as we did for the EEG data. The dashed vertical and solid horizontal lines denote dot onset and chance values, respectively. Shading shows 95% confidence intervals. Bayes factors (BF_10_) quantify the evidence for classification accuracy above chance (*N* = 25). The horizontal line marks a BF_10_ of 1, filled circles show BF_10_ with sufficient evidence for interpretation (in this case showing evidence for the null). See Figure 4 within the main text to compare these results to that yielded from EEG.

## Acknowledgements

We would like to acknowledge that the research conducted for this study took place on Wallumattagal land. This work was supported by an Australian Research Council (ARC) Discovery Project grant awarded to ANR and AW (DP220101067). ANR is supported by an ARC Future Fellowship (FT230100119). AW was supported by MRC (U.K) intramural funding SUAG/093/G116768. We thank Kendall Stead, Brooklyn Gordon, and Kayla Rail for their assistance during data collection.

## Competing Interests

The authors declare no competing interests.

## Author Contributions

BGL, AW, and ANR designed the experiment. SS collected pilot data. BGL, AW, SS, and ANR pre-registered the study. BGL analysed the data. BGL and ANR wrote the paper. AW and SS provided edits.

## Notes

### Competing Interest Statement

The authors have declared no competing interest.

